# Lip movements and lexical features improve speech tracking differently for clear and multi-speaker speech

**DOI:** 10.1101/2023.05.15.540818

**Authors:** Chandra Leon Haider, Hyojin Park, Anne Hauswald, Nathan Weisz

**Affiliations:** University of Salzburg; University of Birmingham

## Abstract

Visual speech plays a powerful role in facilitating auditory speech processing and has been a publicly noticed topic with the wide usage of face masks during the Covid-19 pandemic. In a previous magnetoencephalography (MEG) study we showed that occluding the mouth area significantly impairs neural speech tracking. To rule out the possibility that this deterioration is due to degraded sound quality, in the present follow-up study, we presented participants with audiovisual (AV) and audio-only (A) speech. We further independently manipulated the trials by adding a face mask and a distractor speaker. Our results clearly show that face masks only affect speech tracking in AV conditions, not in A conditions. This shows that face masks indeed primarily impact speech processing by blocking visual speech and not by acoustic degradation. Furthermore, we observe differences in the speech features that are used for visual speech processing. On the one hand, processing in clear speech, but not in noisy speech, is profiting more from lexical unit features (phonemes and word onsets) hinting at improved phoneme discrimination. On the other hand, we observe an improvement in speech tracking driven by the modulations of the lip area in clear speech and conditions with a distractor speaker, which might aid by providing temporal cues for subsequent auditory processing. With this work, we highlight the effects of face masks in AV speech tracking and show two separate ways how visual speech might support successful speech processing.

## 1. Introduction

As the production of an acoustic speech stream is inevitably accompanied by visible mouth movements in face-to-face speech it is unsurprising that the brain has developed mechanisms to interpret both the visual and the auditory input. More importantly, it is also able to integrate both to form a coherent experience. Despite speech processing being dominated by the auditory modality, with the exception of sign language, visual speech plays an important role in the discrimination of neighbouring phonemes as demonstrated by the classic McGurk Effect (Mcgurk & Macdonald, 1976) and improves speech comprehension compared to audio-only settings (Sumby & Pollack, 1954). Research in this field has been brought to general attention with the prevalence of face masks during the Covid-19 pandemic and their impact on speech comprehension in everyday situations. But how does the brain use visual information for speech processing when listening situations are more or less challenging? And how is speech processing affected in such situations, when listeners are deprived of relevant visual input?

Theories about audiovisual speech integration propose two different paths of speech processing enhancements through congruent visual speech (Peelle & Sommers, 2015; Van Engen et al., 2022). On the one hand, visual speech acts as a temporal facilitator for the attentional focus on the acoustic information as visual information precedes the acoustic one (Bourguignon et al., 2020; Chandrasekaran et al., 2009). On the other hand, there is direct information from the mouth and lip area providing information about the phonetic content of speech (e.g. place of articulation) which helps with phoneme discrimination.

In order to investigate these processes on a neural level, new research possibilities have emerged which make it possible to investigate the processing of natural continuous speech presentations. Different approaches have been developed that quantify how “faithfully“ neural responses track speech features (Crosse, Di Liberto, Bednar, et al., 2016; Hauswald et al., 2018; Park et al., 2016). In general terms, these measures of neural speech tracking, implement some association measure between the input signal (e.g., speech envelope) and the output signal (e.g. M/EEG data). The outcome is held to represent the degree to which the respective speech feature (e.g., envelope) is encoded in the brain (Brodbeck et al., 2022; Crosse, Di Liberto, Bednar, et al., 2016).

In the domain of audiovisual speech, past research has revealed that the decoding of the speech envelope from brain data already improves when presenting audiovisual speech in comparison to audio-only speech when the audio input is clear (Crosse et al., 2015) but this visual enhancement further increases when the acoustic information is noisy (Crosse, Di Liberto, & Lalor, 2016). We investigated this phenomenon in the context of face masks as commonly used during the Covid-19 pandemic (Haider et al., 2022). In that continuous audio-visual (AV) speech study, we manipulated the speech signal by adding a face mask and/or a distractor speaker (2x2 Design). For that study, we explored the features of two categories. First, we looked at acoustic features (i.e. speech envelope, pitch and formants) and second, features of lexical segmentation (i.e. phoneme and word onsets) extracted from the speech stimuli. Interestingly, results differed for tracking of acoustic features and lexical segmentation features. On the one hand, tracking of acoustic features was generally affected by masking, while on the other hand tracking of lexical segmentation features was mostly only affected by masking when a distractor was added as well.

However, certain limitations had to be taken into consideration. First, we could not differentiate if the effects on neural tracking via adding a (surgical) face mask were introduced by acoustic distortion of the speech input or by blocking visual speech. Secondly, we only investigated backward modelling (i.e. stimulus reconstruction) which does not allow to link certain experimental effects to specific regions but gives an overall estimate of stimulus representation in the brain. Additionally, the study is limited in its ability to compare the relative contributions of different feature classes, such as acoustic and lexical segmentation, in audiovisual speech processing. A study from our research group using a speech encoding model on the same data as Haider et al. (2022), showed an increased tracking of lip movement in two-speaker speech compared to single-target speaker speech (Reisinger et al., 2023). This might be indicative of a greater usage of lip movements for focussing attention on the succeeding acoustic information as proposed by Van Engen et al. (2022). Another study found increased audiovisual benefit in noisy conditions compared to clear speech conditions through the addition of lexical features in the form of phonemes by using encoding models and canonical correlation analysis (CCA) (O’Sullivan et al., 2021). That study however relied on two separate experiments (one for clear speech and one for noisy speech) which does not allow for a within-subject evaluation of the effect. To this date, it remains unclear on which characteristics (basic lip movements vs. lexical unit features) the enhancement of speech tracking depends on and if the contributions of visual speech can be separated into different functional mechanisms.

With this current study, we want to rule out the more trivial possibility that the negative effect of the face mask is the product of degraded acoustics. Further, we want to separate visual speech benefits generated by basic lip movements and by lexical/phonetic features. For this purpose, we adjusted the experimental design to also include conditions with an audio-only presentation. Now, we can investigate whether the effects of face masks are generated by acoustic distortions or by masked visual cues. An illustration of the new study design can be seen in Figure 1A. In order to investigate how different brain regions contribute to the found effects, we complemented the backward decoding approach with a forward encoding approach in this study thereby linking the effects to MEG channels. Additionally, we are now able to compare models with features of different classes directly. We computed forward models trained on acoustic (spectrogram) features and assessed the impact of adding visual (i.e., lip area) and lexical unit features (i.e., phonemes and word onsets) on speech tracking in clear and noisy conditions. Hereby, we can evaluate how adding complexity to the model improves outcomes and are able to quantify the visual benefit obtained through these additional features.

**Figure 1.**
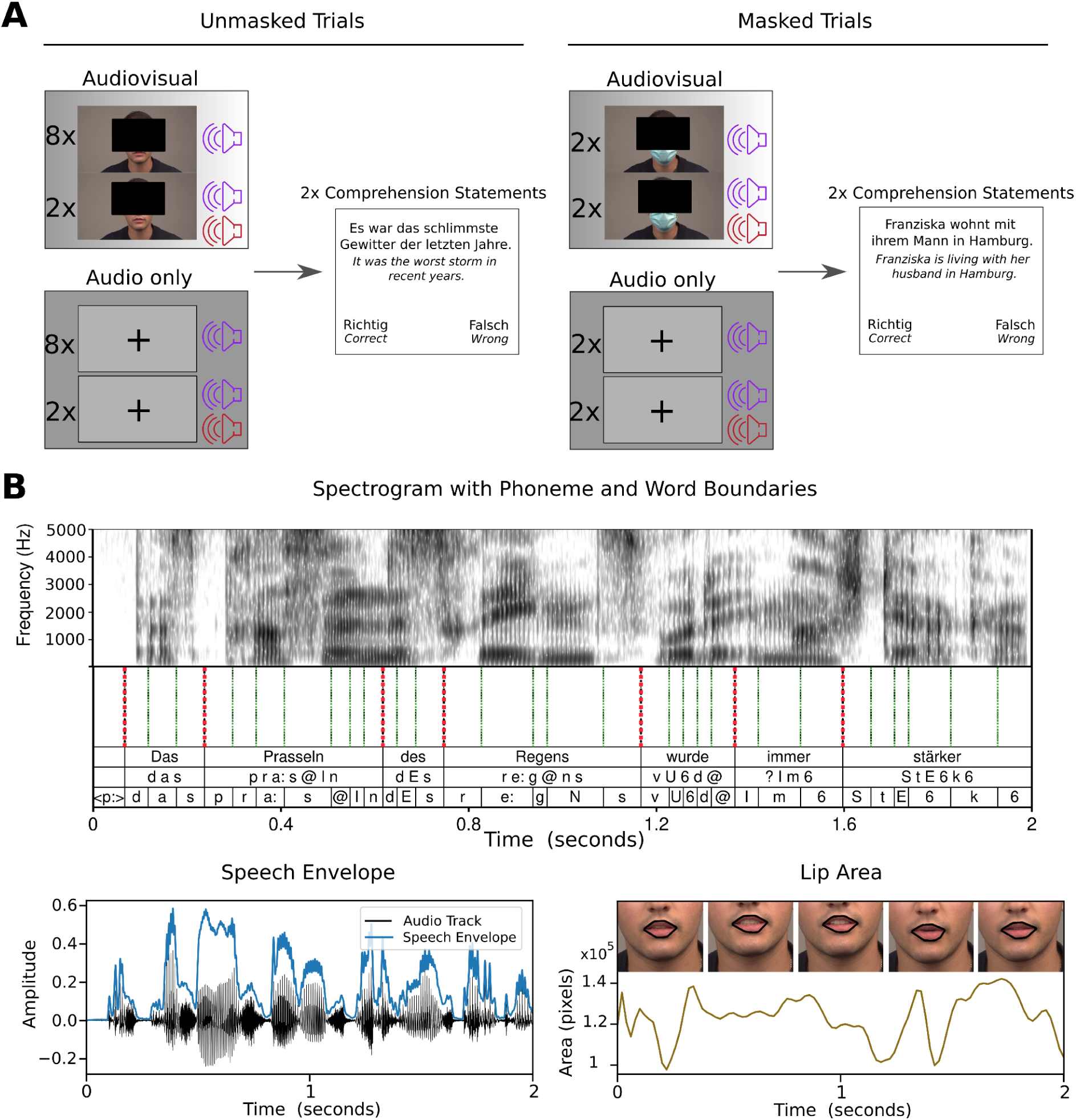
Experimental procedure and speech features. **A** shows the condition design for the experiment with the male speaker as an example. Every participant was presented with 30 trials of ∼ 1 minute each. We conducted audiovisual (AV) (14 trials in total), audio-only (A) (14 trials in total) and visual-only (V) trials (2 trials in total; V trials not depicted in figure). For A and AV trials, we used 10 unmasked trials each. In 8 of these trials, only a single target speaker was presented. In 2 of those trials, we added a second same-sex (distractor) speaker (denoted by the second sound icon). We used 4 masked trials for A and AV speech respectively. In 2 of those trials again only a single target speaker was presented and in 2 trials we added a distractor speaker. After each A and AV trial, we prompted two ‘true or false’ statements to measure comprehension and to keep participants focused. Participants answered via button press (left or right button). **B** shows the investigated speech features. The spectrogram is shown alongside the investigated speech units, phonemes and words underneath (top row: orthographic word; mid row: phonetic word; bottom row: phoneme). The speech envelope can be seen on the bottom left of the figure. On the bottom right of the figure, the extracted lip area can be seen with the presentation of the mouth outline corresponding to the 1st, 13th, 25th, 38th and 50th frames (i.e. 0s, 0.5s, 1s, 1.5s and 2s). All depictions are based on the same two-second-long speech interval. The faces are obscured due to *biorxiv* guidelines.

## 2. Methods

### 2.1. Participants

28 German native speakers (14 female) aged between 20 and 36 years (M = 28.18, SD = 3.79) took part in our study. As we wanted to extend the findings of our previous study, we opted for a similar sample size as in Haider et al. (2022). The exclusion criteria were non-removable magnetic objects, as well as a history of psychiatric or neurological conditions. Recruitment was done via social media and university lectures. All participants signed an informed consent form and were compensated with €10 per hour or course credit. The experimental protocol was approved by the ethics committee of the University of Salzburg and was carried out in accordance with the Declaration of Helsinki.

### 2.2. Stimuli

For this study, we used the same stimulus material as in Haider et al. (2022) but with a reduced number of trials. We used excerpts from four different stories for our recording read out in German. ‘Die Schokoladenvilla - Zeit des Schicksals. Die Vorgeschichte zu Band 3’ (“The Chocolate Mansion, The Legacy” – a prequel of Volume 3”) by Maria Nikolai and ‘Die Federn des Windes’ (“The feathers of the wind”) by Manuel Timm were read out by a female speaker. ‘Das Gestüt am See. Charlottes großer Traum’ (“The stud farm by the lake. Charlotte’s great dream”) by Paula Mattis and ‘Gegen den Willen der Väter’ (“Against the will of their fathers”) by Klaus Tiberius Schmidt were read out by a male speaker.

Stimuli were recorded using a Sony FS100 camera with a sampling rate of 25 Hz for video and a Rode NTG 2 microphone with a sampling rate of 48 kHz for audio. We aimed at a duration for each story of approximately ten minutes, which were cut into ten videos of around one minute each (range: 56–76 s, M = 63 s, SD = 5.0 s). All stories were recorded twice, once without the speaker wearing a surgical face mask and once with the speaker wearing a surgical face mask (Type IIR, three-layer single-use medical face mask, see Fig. 1A). After cutting, all videos were approximately one minute in length (80 AV recordings in total). Thirty of those were presented to each participant (15 with a female speaker, 15 with a male speaker) in order to rule out the sex-specific effects of the stimulus material. The audio track was extracted and stored separately. The audio files were then normalised using the Python function ‘ffmpeg-normalise’. Pre-recorded audiobooks read out by different speakers (one female, one male) were used for the distractor speaker and normalised using the same method. The syllable rate was analysed using a Praat script (Boersma and Weenink, 2001; de Jong and Wempe, 2009). The target speakers’ syllable rates in the 30 trials varied between 3.65 Hz and 4.57 Hz (M = 4.04 Hz, SD = .25). Target and distractor stimuli were all played to the participant at the same volume, which was individually set to a comfortable level at the start of the experiment by using an example audiobook with the target female speaker.

### 2.3. Experimental procedure

Before the start of the experiment, we performed a standard clinical audiometry using an AS608 Basic (Interacoustics, Middelfart, Denmark) in order to assess participants’ individual hearing ability (see Supplementary Material, Figure 1). Out of all participants, four participants showed hearing impairment in higher frequencies (>= 3000 Hz), but reported no to mild subjective impairment. Afterwards, participants were prepared for MEG (see Data acquisition).

We started the MEG measurement with five minutes of resting-state activity (not included in this manuscript). We then adjusted the stimulation volume by presenting an exemplary audiobook and adjusted the volume until participants stated that it was clearly audible and comfortable. One block consisted of six trials of ∼ 1 min length. The condition design is depicted in Figure 1. As an extension to Haider et al. (2022), we added audio-only modality (A) and visual-only modality (V) conditions in addition to the audiovisual modality conditions (AV). For the A trials, we used the same stimuli as the AV trials but did not show the corresponding video and instead presented a fixation cross. This results in 14 AV, 14 A and two V trials. For A and AV conditions respectively, we still wanted to keep the 2x2 design from Haider et al. (2022). Therefore, 8 trials per modality consisted of clear speech (i.e. no mask and no distractor), while in the trials speakers wore a face mask, a second same-sex speaker was added or both (i.e. two trials per one of these conditions). All conditions had an equal amount of female and male speaker trials. In conditions with a second speaker (distractor), the second speaker only started five seconds after the first (target) speaker in order to give participants time to focus on the to-be-attended speaker. Within the first 4 blocks, the story presentation followed a consistent storyline across trials. The fifth block was a mixed speaker block (both female and male speakers) containing continuations from the story ‘Gegen den Willen der Väter’ (male speaker) and ‘Die Federn des Windes’ (female speaker) (three trials each). Some participants also performed six extra visual-only trials which resulted in an additional sixth block. For this, three trials from the story ‘Die Schokoladenvilla - Zeit des Schicksals. Die Vorgeschichte zu Band 3’ and three trials from ‘Das Gestüt am See. Charlottes großer Traum’ were used. As not all participants participated in this sixth block, this was not included in the analysis. With the exception of the V only trials, two unstandardised ‘true or false’ statements regarding semantic content were asked at the end of trials to assess comprehension performance and keep participants focused (Fig. 1A). Additionally, participants rated subjective difficulty twice per condition on a five-point Likert scale for A and AV trials. The participants’ answers were given via button presses. The condition design was shuffled across trials in order to rule out any influence of the specific stimuli material on the results. As an exception to this, we always assigned a trial without a mask and without a distractor as the starting trial of each block in order to let participants adapt to a (possible) new target speaker. Videos were back-projected on a translucent screen with a screen diagonal of 74 cm via a Propixx DLP projector (Vpixx technologies, Canada) ∼ 110 cm in front of the participants. It was projected with a refresh rate of 120 Hz and a resolution of 1920 × 1080 pixels. Including preparation, the experiment took about 2 h per participant. The experiment was coded and conducted with the Psychtoolbox-3 (Brainard, 1997; Kleiner et al., 2007; Pelli, 1997) with an additional class-based library (‘Objective Psychophysics Toolbox’, o_ptb) on top of it (Hartmann and Weisz, 2020).

### 2.4. Data acquisition

We recorded brain data with a sampling rate of 1 kHz at 306-channels (204 first-order planar gradiometers and 102 magnetometers) with a Triux MEG system (MEGIN, Helsinki, Finnland). The acquisition was performed in a magnetically shielded room (AK3B, Vacuumschmelze, Hanau, Germany). Online bandpass filtering was performed from 0.1 Hz to 330 Hz. Prior to the acquisition, cardinal head points (nasion and pre-auricular points) were digitised with a Polhemus FASTRAK Digitizer (Polhemus, Colchester, Vermont, USA) along with around 300 points on the scalp in order to assess individual head shapes. Using a signal space separation algorithm provided by the MEG manufacturer (Maxfilter, version 2.2.15), we filtered noise resulting from sources outside the head and realigned the data to a standard head position, which was measured at the beginning of each block.

### 2.5. Speech feature extraction

All the speech features investigated are depicted in Fig. 1B. The speech envelope was extracted using the Chimera toolbox. By using the default options, the speech signal was filtered forward and in reverse with a 4th order Butterworth bandpass filter at nine different frequency bands equidistantly spaced on the cochlear map between 100 and 10000 Hz (Smith et al., 2002). Then, a Hilbert transformation was performed to extract the envelopes from the resulting signals. These nine bands were then summed up to one general speech envelope and normalised.

The spectrogram was computed similarly. We only adjusted the lower end of the frequency range to 50 Hz, as this then also includes the speech pitch / fundamental frequency (especially for the male speaker). In the end, the resulting nine bands were not summed up but used together as the speech spectrogram.

Phonemes and word onset values were generated using forced alignment with MAUS web services (Kisler et al., 2017; Schiel, 1999) in order to obtain a measure for speech segmentation. We generated two time-series at 50 Hz with binary values indicating an onset of a phoneme or word, respectively.

In order to use these binary features for backward modelling we smoothed the time-series of binary values using a Gaussian window with a width of 20 ms, as 20 ms is the smallest possible window at a sampling rate of 50 Hz. Additionally, we did not distinguish between different phonemes in the case of backward modelling in order to generate a single time series for reconstruction (see Haider et al. (2022). These last two steps were skipped when features were analysed using forward models because in this case binary features can be investigated and a multivariate input (i.e. different individual phonemes) is beneficial. In this case, we ended up with 63 time series, one for each phoneme, containing only zeros and ones.

The lip area was extracted from the video recordings using a Matlab function by Park et al. (2016). Through this, we generated a time series of the lip area in pixels with a sampling rate of 25 Hz (i.e. the frame rate of the video recording).

In the end, all features were sampled to 50 Hz to match the sampling rate of the corresponding brain signal, as most speech-relevant signals present themselves below 25 Hz (Crosse et al., 2021).

### 2.6. MEG preprocessing

The raw data was analysed using Matlab R2020b (The MathWorks, Natick, Massachusetts, USA) and the FieldTrip toolbox (Oostenveld et al., 2011). As part of our standard pipeline, we first computed 50 independent components to remove eye and heart artefacts. We removed on average 2.29 components per participant (SD = .86). We further filtered the data using a sixth-order zero-phase Butterworth bandpass filter between 0.1 and 25 Hz. Finally, we downsampled our data to 50 Hz for more efficient computation while still preserving sufficient information from our data (Crosse et al., 2021).

### 2.7 TRF model fitting

In order to prepare the data for fitting the TRF models, we first z-scored the MEG data of all 306 channels and rescaled the speech envelope and lip area to values between 0 and 1. This rescaling was preferred over z-scoring, as this does not introduce negative values into only positive sign features. We used the mTRF Toolbox (Crosse, Di Liberto, Bednar, et al., 2016) to reconstruct stimulus features (in the case of backward modelling) or to predict brain data (in the case of forward models). This was done similarly to Haider et al. (2022). For a more detailed discussion about this method please refer to Crosse et al. (2016) and Crosse et al. (2021). We trained two separate models on audiovisual (AV) clear speech (i.e. AV speech without a face mask and without a distractor) and on audio-only (A) clear speech respectively. We then used these models to reconstruct the stimulus feature or predict brain activity for their respective modality (AV model for AV conditions and A model for A conditions) but across other conditions (i.e. for trials with masks and with distractors). As the second distractor speaker only starts after 5 seconds into the trial, we removed the first 5 seconds of each distractor trial from the analysis. For both forward and backward modelling, we used time lags from -50 to 500 ms to train our models. Regularisation parameter (lambda) between 10^-6^ … 10^6^ was determined using a seven-fold leave-one-out cross-validation (Willmore & Smyth, 2003).

In order to completely rule out the possibility that a single outlier trial influences the model training, we again made use of a leave-one-out procedure. We used seven clear speech trials for model training and the eighth clear speech trial for model testing. We then looped across the trials so that every trial was once used for testing and otherwise used for training. In the end, we averaged the Fisher-z-transformed values across each iteration together. Overall, we used ∼ 7 minutes for training our A and AV models and ∼ 1 of testing for the clear speech condition in each iteration. For all other three conditions (i.e. No Mask + Distractor; Mask + No Distractor; Mask + Distractor), we used all trials for testing which was ∼ 2 minutes each.

#### 2.7.1 Backward modelling

With this approach we are able to map the brain response back to the stimulus feature (e.g. the speech envelope) in order to acquire a measure of how well a certain feature is encoded in the brain. This model has the benefit that the whole brain activity is taken into account when reconstructing a feature, which holds information about a general representation of a certain stimulus feature in the brain. By this approach, you obtain a single correlation value (Pearson’s r) as the measure of how well a feature can be reconstructed, which makes it easily interpretable. However, it is not possible to reconstruct multidimensional features (e.g. a speech spectrogram) but only one-dimensional features (e.g. the speech envelope). Furthermore, by using this approach spatial information on which brain regions are involved in the processing of certain stimulus characteristics is lost.

#### 2.7.2 Forward modelling

By using this approach, we are mapping the stimulus (-feature) ‘forward’ to the individual MEG channels. We therefore acquire a measure of how well a certain stimulus feature is encoded in the individual channels. In contrast to backward models, this model cannot give a measure of how well a feature is generally encoded in the whole brain, as it does not account for intercorrelations between the channels. However, as we get a prediction accuracy measure for each individual channel, it is possible to acquire a spatial distribution in the brain and test hypotheses about the contribution of different brain regions. Additionally, we can use multivariate inputs in order to predict brain data (e.g. spectrogram) which allows us to evaluate more complex (multidimensional) features.

### 2.8 Statistical analysis

For the analysis of backward models and separately for the A and the AV conditions, we performed 2x2 repeated measures ANOVA with the factors *Mask* (no face mask vs. face mask) and *Distractor* (no distractor speaker vs. distractor speaker) and the obtained Fisher z-transformed correlation coefficients (i.e. reconstruction accuracy) as dependent variables. For evaluating the effects of *Modality* (A vs. AV) we additionally performed a 2x2x2 repeated measures ANOVA with the within-factors Modality, Mask and Distractor.

For the behavioural results (comprehension performance and subjective difficulty), we also used repeated measures ANOVA with the same factors Modality, Mask and Distractor. We used comprehension performance scores (i.e. the percentage of correct answers) and averaged subjective difficulty ratings respectively as dependent variables. For one participant behavioural data was missing resulting in 27 subjects for this analysis.

We analysed the forward models with one-tailed dependent sample cluster-based permutation tests with 10000 permutations (Maris & Oostenveld, 2007). We used the *maxsum* method implemented in Fieldtrip for the calculation of the t-statistic. We furthermore investigated the effect size of each cluster by averaging the Cohen’s d values across all significant channels in that cluster. A one-tailed test was chosen because we tested for differences in simple versus additive models, whose performance must be at least on the level of the simple model. We restricted our analysis to combined gradiometers, as these allow for a better spatial resolution. For that, the correlation coefficients of each gradiometer pair were averaged.

## 3. Results

### 3.1 Behavioural Results

For comprehension performance we computed the percentage of correctly answered questions for each condition for each participant. We analysed the data according to our 2x2x2 design with the factors *Modality* (A vs. AV), *Mask* (no mask vs. mask) and *Distractor* (no distractor vs. distractor). The only significant main effect was for the *Distractor*, revealing a significant decrease in comprehension performance *(F(1,26) = 5.73, p < .024, η_p_² = .18)*. The interaction between *Modality* and *Mask* was also significant, showing only a decrease through the mask of comprehension performance in AV conditions (*F(1,26) = 6.28, p < .019, η_p_² = .20)*. Please note that for most conditions (all except clear speech conditions), we only prompted four comprehension statements per subject with a guess rate of 50%. Therefore, these results should be interpreted with care. All other results were not significant and are shown in the Supplementary Material (Table 1).

The subjective difficulty was calculated by averaging the ratings (as indicated on a five-point Likert scale by the participant) per condition. Results show a significant effect of *Modality (F(1,26) = 11.84 p = .002, η_p_² = .31), Mask (F(1,26) = 8.71 p = .007, η_p_² = .25)* and *Distractor (F(1,26) = 109.68, p < .001, η_p_² = .81)*. The interactions between *Modality* and *Mask* were also significant (*F(1,26) = 7.23 p = .012, η_p_² = .22)* as well as between *Modality* and *Distractor* (*F(1,26) = 5.04 p = .034, η_p_² = .16)*. Participants reported more difficulty in face mask trials than in no face mask conditions only in the AV condition, while it stayed the same for the A conditions. Moreover, AV stimuli had a buffering effect on the influence of the distractor by reducing its impact as indicated by the significant interaction between *Modality* and *Distractor*. All other results were not significant and are shown in the Supplementary Material (Table 1).

### 3.2 Backward modelling is only affected by face masks in audiovisual conditions not in audio-only conditions

By using the mTRF Toolbox in the backward direction (Crosse, Di Liberto, Bednar, et al., 2016), we calculated one correlation coefficient per condition and per subject for each investigated feature. The features used here were the speech envelope (as an acoustic feature), phoneme and word onsets (as lexical segmentation features) and lip area (as a visual feature). The correlation coefficients were then Fisher-z transformed and averaged across each condition.

For statistical analysis, we used two 2x2 ANOVA models separately for A and AV conditions and performed a 2x2x2 ANOVA to investigate the influence of *Modality* and the interactions. In general, all reconstruction accuracies investigated were heavily influenced by the addition of a distractor speaker. Therefore, the main effect of *Distractor* will not be reported here, but is visually depicted in Figure 2 and in Supplementary Material (Table 2 - 4).

**Figure 2.**
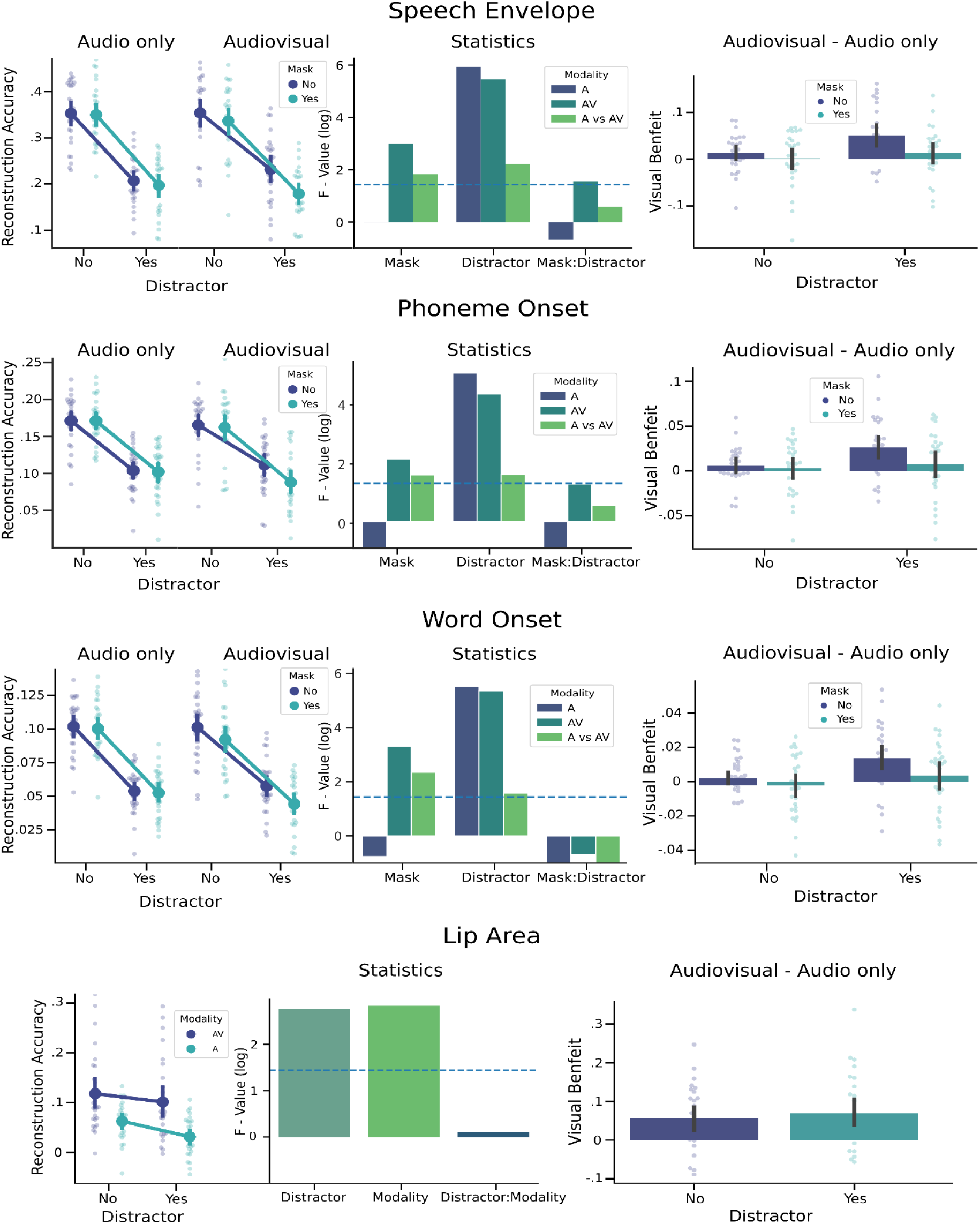
Visual illustration of the results of the backward modelling. Visual illustration of the results of the backward modelling for the investigated features speech envelope, phoneme onset, word onset and lip area. On the left side, the raw reconstruction accuracy is depicted split up for audio-only (A) and audiovisual (AV) conditions, except for lip area. Next to it on the right, the statistical evaluation of the effects of *Mask* and *Distractor* and their interaction are shown with log-scaled F-values. The dashed horizontal blue line represents the critical F-value (α = 5%) of 4.20 (1.44 in log space). The light green bar represents the interaction with *Modality* (i.e. the difference between A and AV) for each effect. On the right hand side, the visual benefit for each subject is shown (i.e. AV - A). For the lip area feature, it was not possible to evaluate the effects of the face mask, therefore only the effects of Distractor and Modality can be seen. All error bars in the figure represent 95% confidence intervals calculated by *seaborn*.

The visual benefit was calculated by subtracting the reconstruction accuracies in the A conditions from the AV conditions. Results are shown in Figure 2. For the conditions when the speakers wore a face mask, we could not compute the lip area. However, we investigated the lip area tracking in the AV conditions and also the tracking of the unseen lip area in the A condition.

#### 3.2.1 Speech Envelope

In AV conditions, the reconstruction of the speech envelope showed impairment through a face mask (*F(1,27) = 20.15, p < .001 η_p_² = .42*) as well as an interaction with a distractor (*F(1,27) = 4.79, p = .037 η_p_² = .15)*. For the A conditions both these effects are missing (*Mask: F(1,27) = 0.999, p = .327 η_p_² = .04; Mask:Distractor: F(1,27) = 0.51, p = .48, η_p_² = .02)*. The effect of modality shows a significant main effect (*F(1,27) = 10.42, p = .003, η_p_² = .28*), as well as a significant interaction with *Mask* (*F(1,27) = 6.20, p = .019, η_p_² = .19*) and *Distractor* (*F(1,27) = 9.24, p = .005 η_p_² = .26)*. The threefold interaction (*Modality:Mask:Distractor*) however is not significant (*F(1,27) = 1.82, p = .189, η_p_² = .06*).

Investigating the confidence intervals for each condition individually reveals that only for the distractor condition without a face mask the visual benefit is significantly above zero (*95% CI [.0295, .0717]*). Hereby we can confirm the idea that visual speech has a buffering effect on tracking of the speech envelope especially in conditions where the audio input is unclear. Face masks only affect tracking in AV conditions, but do not show any significant impact on tracking in A conditions. Similar results can be seen for the lexical segmentation features of phoneme onset and word onset.

#### 3.2.2 Phoneme Onset and Word Onset

In AV conditions, the reconstruction of the phoneme onsets showed impairment through a face mask (*F(1,27) = 10.49, p < .001 η_p_² = .003*) and a trend for the interaction with the distractor condition (*F(1,27) = 4.07, p = .054 η_p_² = .13)*. For the A conditions both these effects are not significant (*Mask: F(1,27) = 0.06, p = .809 η_p_² = .00; Mask:Distractor: F(1,27) = 0.04, p = .843, η_p_² = .00)*. The effect of modality shows a significant main effect (*F(1,27) = 9.03, p = .005, η_p_² = .25*), as well as a significant interaction with *Mask* (*F(1,27) = 5.75, p = .024, η_p_² = .18*) and *Distractor* (*F(1,27) = 5.84, p = .023, η_p_² = .18).* The threefold interaction (*Modality:Mask:Distractor*) however is not significant (*F(1,27) = 1.84, p = .186, η_p_² = .06*). Again the only condition with significant visual benefit was the distractor condition without a face mask (*95% CI [.0147, .0381]*).

In AV conditions, the reconstruction of the word onsets showed impairment through a face mask (*F(1,27) = 27.15, p < .001, η_p_² = .50*) but no interaction with the distractor condition (*F(1,27) = .50, p = .487, η_p_² = .02)*. For the A conditions both these effects are again not significant (*Mask: F(1,27) = .50 p = .499, η_p_² = .02; Mask:Distractor: F(1,27) = 0.00, p = .978, η_p_² = .00)*. The effect of modality shows a significant main effect (*F(1,27) = 8.57, p = .007, η_p_² = .24*), as well as a significant interaction with *Mask* (*F(1,27) = 10.44, p = .003, η_p_² = .28*) and *Distractor* (*F(1,27) = 4.88, p = .036, η_p_² = .15).* As for the other features, the threefold interaction (*Modality:Mask:Distractor*) is not significant (*F(1,27) = .26, p = .616, η_p_² = .01*). In this case visual benefit is significant in both conditions without a mask, i.e. without a distractor (*95% CI [.0007, .0079]*) and with a distractor (*95% CI [.0068, .0208]*).

For both lexical segmentation features, results are similar to the effects observed for tracking of the speech envelope. Again, face masks only have an impact in AV conditions, not in A. The visual benefit is largest in the condition with a distractor speaker but only when the speaker does not wear a face mask. For the tracking of the word onsets, both conditions without a face mask show significant visual benefit (despite being much smaller in the clear speech conditions).

#### 3.2.3 Lip Area

As we could not compute the lip area for the trials in which speakers wore a face mask (as the mouth is occluded in the video), we computed a 2x2 ANOVA with the factors Modality (A vs AV) and Distractor (no distractor vs distractor). Both main effects are significant (Modality: *F(1,27) = 16.90, p = <.001, η_p_² = .39; Distractor: F(1,27) = 15.86, p = <.001, η_p_² = .37*), while the interaction was not significant (*F(1,27) = 1.12, p = .299, η_p_² = .04*). As expected, both conditions show above zero visual benefit (*95% CI*, no distractor: *[.0255, .0852]*, distractor: *[.0356, .1041]*). These results highlight the importance of controlling for effects generated by correlated audio features when investigating visual speech in an AV paradigm. This is further supported by strong correlation between audio only lip area tracking values and audio only speech envelope tracking values (no distractor: r(26) = .605, p <.001, distractor: r(26) = .472, p = .01).

### 3.3 Forward modelling reveals improved tracking across conditions through modulations of the lip area and increased visual benefit through lexical features only in clear speech

By using the mTRF Toolbox in the forward direction (Crosse, Di Liberto, Bednar, et al., 2016), we calculated one correlation coefficient per condition, per channel and per subject for each investigated feature. In contrast to backward modelling, we can now investigate features of multidimensional scale, such as the speech spectrogram or individual phonemes. As speech features are proven to be strongly intercorrelated, the use of additive models is highly beneficial to investigate differences and potential gains through adding additional features to the models. For this analysis we computed a (base) acoustic model (spectrogram), a spectrogram + lip model (spectrogram + lip area) and a spectrogram + lexical model (spectrogram + phonemes + word onsets). The resulting correlation coefficients from the models were Fisher-z transformed and averaged within conditions. As we have shown for the backward models, the effect of the mask is only affecting features in conditions with visual input significantly. Therefore, we did not include conditions with masks in this following result section, but instead focused on the visual benefit in clear speech and speech with a second (distractor) speaker. To investigate the effects statistically, we used a one-tailed dependent-sample cluster-based permutations test, as described in the Methods section.

#### 3.3.1 Spectrogram Model

We used the spectrogram model as our baseline model, shown in Figure 3. For this model, we only estimated statistical contrasts between the A and AV speech for clear speech and distractor speech. We only found an increased tracking in AV compared to A speech in the distractor condition for the right temporal and occipital sensors (*t_sum_ = 50.27, p = .004, d = 1.14*) and a second cluster in left temporal and frontal regions (t*_sum_* = 37.69, p = .009, d = .79), while we did not find a significant difference for the clear speech condition. This effect corresponds well to the observation of the backward models, which confirm that visual benefit is beneficial in the conditions with an unclear speech acoustic.

**Figure 3.**
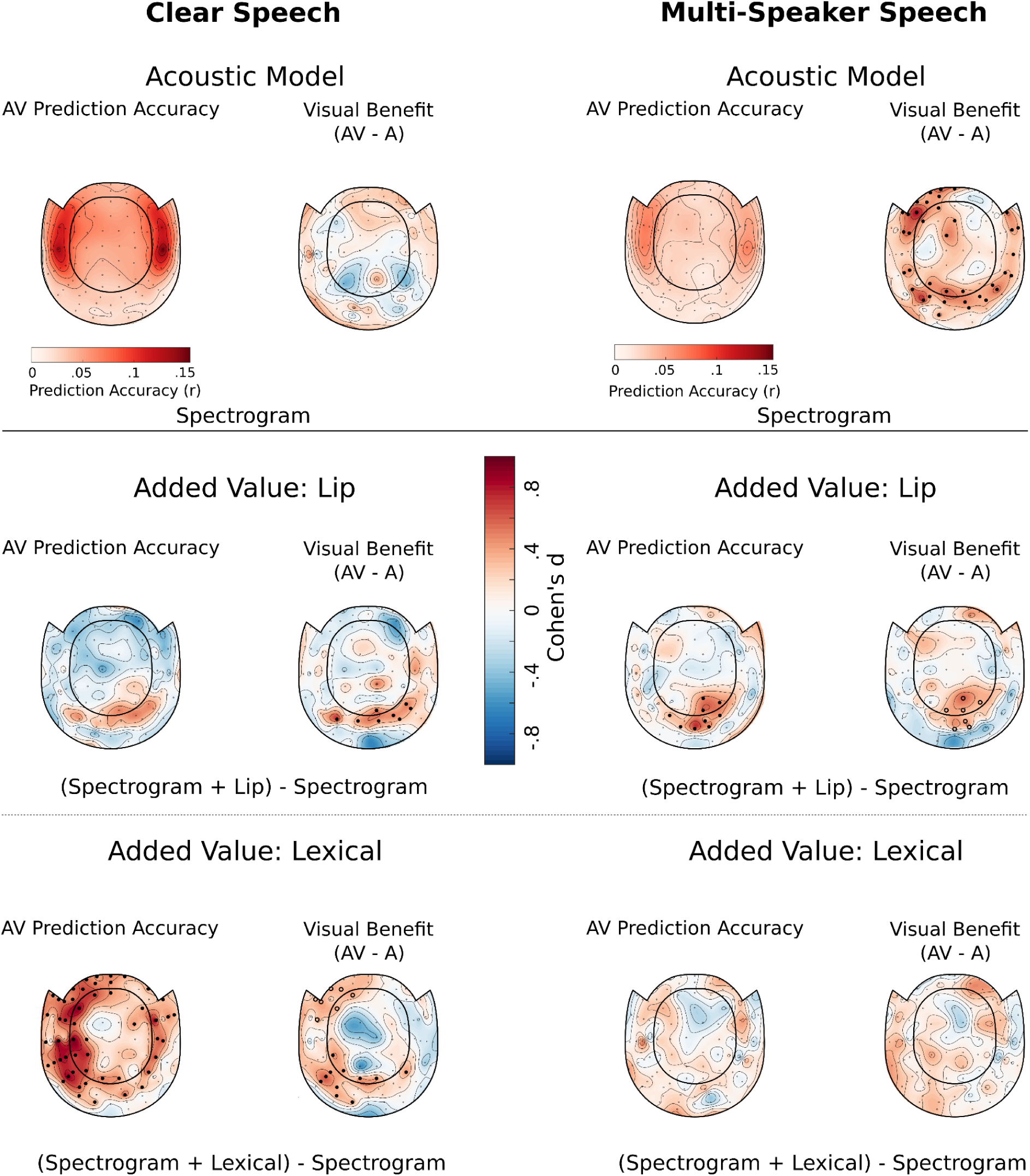
Visual illustration of the results of the forward modelling and added value. Results of our forward modelling analysis. The first row shows the prediction accuracy and visual benefit for all combined gradiometers of the (baseline) acoustic model based on the spectrogram. On the left side, these values were acquired from brain data of participants listening to clear speech; on the right side, they were listening to a multi-speaker (distractor) speech. Statistical analysis for the visual benefit (i.e. AV - A) for the acoustic model was calculated. The second and third rows show the added value of visual features (lip area) and lexical (unit) features (phonemes and word onsets) respectively. Specifically, they represent the difference between the spectrogram model and the additive model (i.e. spectrogram + lip or spectrogram + lexical) for prediction accuracies and visual benefit. This again is done for clear speech and distractor speech. Filled dots represent channels wIth significant differences computed by a one-tailed cluster-based permutation test (α = 5%). Circles represent trends (p < .10).

#### 3.3.2 Added Value: Spectrogram + Lip Model

The additive model of spectrogram and lip area showed no significant improvement in the clear speech condition over the acoustic-only spectrogram model for overall prediction accuracy. However, a significant increase in visual benefit in the clear speech conditions can be observed (t*_sum_* = 19.42, p = .030, d = .69). Moreover, there is an increase in prediction accuracy in the distractor speech condition (*t_sum_ = 21.68, p = .031, d = .67*) over occipital areas. Additionally, this model shows a trend for increased visual benefit over the spectrogram-only model in occipital areas (*t_sum_ = 14.37, p = .067, d = .57*) in distractor conditions. The added value of the lip area seems not to be restricted to either clear speech or distractor speech and improves speech tracking in general.

#### 3.3.3 Added Value: Spectrogram + Lexical Model

In contrast to the Spectrogram + Lip model, the increase through the Spectrogram + Lexical model shows increases restricted to the clear speech condition. On the one hand, we observe a widespread increase in prediction accuracy in the clear speech condition (*t_sum_ = 150.40, p < .001, d = 1.08*). Furthermore, the increase in visual benefit in the clear speech condition over the acoustic spectrogram model can be observed significantly in visual areas (t*_sum_* = 24.01, p = .013, d= .63) and in an additional trend in left frontal areas (t*_sum_* = 13.16, p = .078, d = .47). On the other hand, we do not see any effect in the distractor condition neither for prediction accuracy nor for visual benefit. This is in contrast to the added value of the spectrogram + lip model which seems to generally improve speech tracking across levels of acoustic clarity.

## Discussion

In a previous work, we showed that face masks impair the reconstruction of acoustic features and of features of lexical segmentation in the presence of a distractor speaker (Haider et al., 2022). As we however only investigated the effects of face masks in AV speech, we could not differentiate the contribution of (blocked) visual speech and the distortion of the acoustic signal to this face mask effect. Furthermore, our study only employed a backward modelling approach, which limits our ability to investigate the contribution of speech features to specific brain regions. In this study, we aimed to answer these remaining questions. The backward modelling approach revealed that the negative effects of face masks on reconstruction accuracy for the investigated speech features can be primarily attributed to the absence of visual information, whereas the acoustic distortion by surgical face masks appears to have a negligible impact on speech tracking. We can further add to this, that additional visual information in general has a buffering effect on the decrease in tracking when the audio input becomes unclear which points to an increase in visual contribution in this situation (Crosse, Di Liberto, & Lalor, 2016; Crosse et al., 2015; Golumbic et al., 2013).

With the forward modelling, we can show improved speech tracking of an acoustic feature (i.e., spectrogram) in difficult listening situations through visual speech. We can provide evidence that beyond acoustic tracking, visual benefit is presenting itself in two separate ways: When listening is relatively unchallenging, visual speech improves neural speech tracking through more complex lexical features. When speech becomes more challenging, the brain only relies on tracking the modulations of the speaker’s lip area, possibly to focus attention on the target.

### Face masks negatively affect speech reconstruction, comprehension and subjective difficulty only in an audiovisual not in an audio-only presentation

As hypothesised in our previous study (Haider et al., 2022), the effect face masks have on tracking is only relevant in AV settings. This is shown by significant effects of the face mask for all features in the AV speech condition, with all these effects being absent in the A speech conditions. This supports the idea that despite the changes to the acoustic signal (Corey et al., 2020; Haider et al., 2022; Homans & Vroegop, 2021), the negative impact of face masks is indeed produced by the missing visual speech input. This holds also true for the behavioural results of comprehension performance and participants’ subjective difficulty ratings, which measure the constraints people face more directly. This study again shows the negative effects of face masks on speech processing, as quantified on both neural and behavioural levels. While the negative impact of a decreasing level of speech comprehension is obvious, an increased subjective difficulty has been shown to lead to social withdrawal in some individuals (Hughes et al., 2018).

In comparison to our previous study, where face masks had the most detrimental effects in a distractor condition, the impact was more general in the present study. Especially, the effects for the reconstruction of the word onset do not show the same pattern. In the previous study, we found an interaction of the face mask with the distractor, hinting at a stronger effect of face masks in difficult listening situations, which is missing in the present study. Despite reanalyzing the old data and adjusting the analysis to be identical to this study (i.e. manual resampling of binary feature, smoothing with 20 ms window, adjusting time lags to [-50 500] and most importantly restricting model training to only ∼ 7 minutes), the effect remained stable. It can only be attributed to (uncontrolled) differences in population or by the fact that participants in the former study had fewer experimental conditions, but more trials per condition and could therefore adjust better to each individual listening situation (e.g. listening to a target speaker in multi-speaker speech).

Nonetheless these inconsistencies, our results have important implications for the use of face masks in critical contexts, such as hospitals. Following our findings we would advocate in favour of face masks which allow visual speech still to be processed. As transparent face shields proved to be of little use in stopping the transmission of viruses (Lindsley et al., 2021) and transparent face masks are distorting the acoustic signal severely (Brown et al., 2021; Corey et al., 2020), we cannot make final conclusions here. Using transparent face masks might be a valuable solution in clinical settings when communicating with hearing-impaired individuals (Atcherson et al., 2017; Kratzke et al., 2021).

### Lip movements are a general facilitator of speech tracking, while lexical features only improve speech tracking in clear speech

A past study has already convincingly demonstrated the visual benefit of lip movements in a neural tracking context (Reisinger et al., 2023). However, many studies have only focused on visual tracking in a visual-only speech paradigm. Hauswald et. al (2018) and Bourguignon et al. (2020) demonstrate that tracking of lip movements can be first observed in the visual cortex. This information is then extracted in the right angular gyrus and forwarded to auditory cortices. Here, they are mapped to the predicted corresponding acoustic speech. Although this study focused on separate visual and audio-only speech, the conclusion that visual speech is predicting expected acoustic input to auditory cortices seems nonetheless plausible, because in natural speech the visual information precedes the acoustic one (Bourguignon et al., 2020). Despite an overwhelming amount of behavioural evidence for visual enhancement of speech processing, only little research has been published that demonstrates this on a neural level. Reisinger et al. (2023) showed an increase in the tracking of lip movements when listening to a target speaker while a distractor speaker was also present. They also found that subjects differ substantially in the extent to which they incorporate visual speech for speech processing, a result previously demonstrated by Aller et al. (2022). However, Reisinger et al. (2023) corroborate this finding further by showing that people with this higher reliance on visual speech show worse comprehension performance and higher subjective difficulty ratings in conditions when the mouth area was occluded. In general, our findings show improved visual benefit through lip movements in conditions where audio input is unclear as hypothesised by the concept of ‘inverse effectiveness’ (Meredith & Stein, 1986), but also in clear speech conditions. This benefit is primarily observed in occipital sensors which might fit well with the proposed processing described by Hauswald et al. (2018) and Bourguignon et al. (2020) leading from the visual to the auditory cortex.

Contrasting the effect of lip movements, the addition of higher-order lexical features to the model does not show this pattern. Only in clear speech, lexical information seems to improve tracking in general as well as showing improved visual benefit over the acoustic model. The topography of these effects is also different. Visual benefit through lexical features seem to concentrate on the left hemisphere in mainly temporal regions and may represent access to higher level lexical/phonetic information (Hickok & Poeppel, 2007; Zatorre et al., 1992), although there is evidence that processing of lexical units is occurring already in visual areas (Nidiffer et al., 2021).

We interpret this as a shift in how speech processing is facilitated: While audio speech is clear, visual information is used to extract phonemic information from lip movements and other components of visual speech and integrate it with the acoustic information. In the presence of a distracting speaker, more low-level visual information (e.g., mouth area) can be used to disambiguate the speaker streams. Therefore, the brain is more sensitive to rudimentary modulations of the lip area than to extracting complex lexical information through e.g. place of articulation. In terms of the two processing modes proposed by Peelle & Sommers (2015), this would represent a stronger reliance on early visual facilitation through temporal cues derived from lip movements in distractor speech and also clear speech. In clear speech, the brain additionally benefits from the late process of phoneme discrimination more prominently observed in the left hemisphere. As a past study has shown, stronger neural tracking can be seen as an enhancement of representation of the target over the distractor speech stream (Orf et al., 2022). We therefore argue that in a multi-speaker listening situation, the processing of lip movements improves target-over-distractor representations in this regard, while lexical features do not. The benefit of visual features (i.e., lip area) in the multi-speaker condition might be exclusive to situations with a distractor speaker and might not be generalisable to other signal-to-noise manipulations such as adding white noise.

However, further research is needed as other studies do not show this pattern. Similarly to our study O’Sullivan et al. (2021) investigated audiovisual benefit (i.e. AV > A + V) through lexical features in clear and noisy speech via EEG. In contrast to our study, they found an increase in audiovisual benefit in the noisy condition compared to the clear speech condition for lexical features, while controlling for acoustic features. A few differences to our study have to be taken into account. First, they investigated the AV benefit, while we can only make statements about visual benefits for speech processing. Secondly, two separate experiments with different subjects were conducted in clear and noisy speech respectively which does not allow for a within-subject comparison of effects. Third, they created a difficult listening situation by adding noise compared to adding a second speaker in our study, which might involve different processes.

### Future directions

The impact of face masks on speech tracking can indeed be traced back to missing visual speech. However, we observed mainly young normal hearing subjects in this and our past study, which leaves the question open how older individuals or individuals with hearing loss are affected by face masks.

As previous studies have shown, the interindividual variability for the incorporation of visual information for speech processing is relatively high (Aller et al., 2022; Reisinger et al., 2023). One interesting subject group in that regard are again individuals suffering from hearing impairment because a past study has already shown that these subjects do profit more from added visual information compared to normal hearing subjects (Puschmann et al., 2019). These proposed benefits were however never quantified on a neural level. Moreover, results from this study and the study from O’Sullivan et al. (2021) do not show the same pattern. However, differences in approach might be the reason for this. A future within-subject study which makes it also possible to investigate audiovisual benefit (AV > A + V) in clear and distractor speech might be able to reconcile the two results. Additionally, a systematic investigation on how different degradations of the speech signal (e.g. multi-speaker speech, adding noise or vocoding) affect tracking results might be beneficial in completing the picture of speech tracking in difficult listening situations.

## Conclusion

While some differences could be observed to our previous results, we show here that the negative impact of face masks on neural speech tracking (in this case reconstruction accuracy) can be traced to a blocking of the visual speech of the speaker, while acoustic degradations through (surgical) face masks seem to have minimal impact. Using forward modelling we show that adding features to a baseline acoustic model affects the visual benefits differently according to the framework by Peelle and Sommers (2015): Lip movements improve visual benefit in distractor speech conditions and clear speech conditions hinting at a general process in AV speech tracking as a temporal facilitator. Lexical features show visual benefit only in the clear speech conditions, hinting at a more demanding late processing for phoneme discrimination.

Declaration of Competing Interest Statement

The authors have declared no competing interest.

## Acknowledgements

This work is supported by the Austrian Science Fund P34237 (“Impact of face masks on speech comprehension”). Sound icon made by *Smashicon* from www.flaticon.com.

I would like to thank the whole research team and especially Juliane Schubert for her help as a speaker as well as giving direct and strict but most often correct feedback.

## Code Availability Statement

The code to reproduce statistical analyses can be found at https://gitlab.com/CLH96/visualspeechenhancements. Further code can be supplied upon request.

