## Supplementary Material for "Lip movements and lexical features improve speech tracking differently for clear and multi-speaker speech"

Table 1. 2x2x2 ANOVA for the behavioural measures of *Comprehension* and *Subjective Difficulty* with the factors *Modality, Mask* and *Distractor*

| | **Comprehension** | | | **Subjective Difficulty** | | |
|---|---|---|---|---|---|---|
| | **F(1,26)** | ***p*** | $\eta_p^2$ | **F(1,26)** | ***p*** | $\eta_p^2$ |
| Modality | .08 | .769 | .00 | **11.84** | **.002** | **.31** |
| Mask | 2.30 | .141 | .08 | **8.71** | **.007** | **.25** |
| Distractor | **5.73** | **.024** | **.18** | **109.68** | **<.001** | **.81** |
| Modality: Mask | **6.28** | **.019** | **.19** | **7.23** | **.024** | **.22** |
| Modality: Distractor | 1.41 | .246 | .05 | **5.04** | **.033** | **.16** |
| Mask: Distractor | .06 | .807 | .00 | .01 | .118 | .00 |
| Modality: Mask: Distractor | 1.19 | .286 | .04 | .25 | .186 | .01 |

Table 2. 2x2 ANOVA for the AV Model with factors *Mask* and *Distractor*

| | **Main effect: *Mask*** | | | **Main effect: *Distractor*** | | | **Interaction** | | |
|---|---|---|---|---|---|---|---|---|---|
| | **F(1,27)** | ***p*** | $\eta_p^2$ | **F(1,27)** | ***p*** | $\eta_p^2$ | **F(1,27)** | ***p*** | $\eta_p^2$ |
| Envelope | **20.15** | **<.001** | **.42** | **233.5** | **<.001** | **.90** | **4.79** | **.037** | **.15** |
| Phoneme Onset | **10.49** | **.003** | **.28** | **120.4** | **<.001** | **.82** | 4.07 | .054 | .13 |
| Word Onset | **27.15** | **<.001** | **.50** | **211.58** | **<.001** | **.89** | .50 | .487 | .02 |
| Lip Area | – | – | – | 2.47 | .128 | .08 | – | – | – |

Table 3. 2x2 ANOVA for the A Model with factors *Mask* and *Distractor*

| | **Main effect: *Mask*** | | | **Main effect: *Distractor*** | | | **Interaction** | | |
|---|---|---|---|---|---|---|---|---|---|
| | **F(1,27)** | ***p*** | $\eta_p^2$ | **F(1,27)** | ***p*** | $\eta_p^2$ | **F(1,27)** | ***p*** | $\eta_p^2$ |
| Envelope | .999 | .331 | .04 | **373.92** | **<.001** | **.93** | .51 | .48 | .02 |
| Phoneme Onset | .06 | .809 | .00 | **259.74** | **<.001** | **.91** | .04 | .843 | .00 |
| Word Onset | .50 | .499 | .02 | **251.86** | **<.001** | **.90** | .00 | .978 | .00 |
| Lip Area | – | – | – | **18.486** | **<.001** | **.41** | – | – | – |

Table 4. 2x2x2 ANOVA with factors *Modality*, *Mask* and *Distractor*

| | Speech Envelope | | | Phoneme Onset | | | Word Onset | | | Lip Area | | |
|---|---|---|---|---|---|---|---|---|---|---|---|---|
| | **F(1,27)** | ***p*** | $\eta_p^2$ | **F(1,27)** | ***p*** | $\eta_p^2$ | **F(1,27)** | ***p*** | $\eta_p^2$ | **F(1,27)** | **p** | $\eta_p^2$ |
| Modality | **10.42** | **.003** | **.28** | **9.03** | **.005** | **.25** | **8.57** | **.007** | **.24** | **16.90** | **<.001** | **.39** |
| Mask | **18.06** | **<.001** | **.41** | **5.19** | **.031** | **.16** | **14.35** | **<.001** | **.35** | – | – | – |
| Distractor | **401.44** | **<.001** | **.94** | **250.97** | **<.001** | **.90** | **359.3** | **<.001** | **.93** | **15.86** | **<.001** | **.37** |
| Modality: Mask | **6.20** | **.019** | **.19** | **5.75** | **.024** | **.18** | **10.44** | **.003** | **.28** | – | – | – |
| Modality: Distractor | **9.24** | **.005** | **.26** | **5.84** | **.023** | **.18** | **4.88** | **.036** | **.15** | 1.12 | .299 | .04 |
| Mask: Distractor | **5.78** | **.023** | **.18** | 2.61 | .118 | .09 | 0.30 | .589 | .01 | – | – | – |
| Modality: Mask: Distractor | 1.82 | .189 | .06 | 1.84 | .186 | .06 | 0.26 | .616 | .01 | – | – | – |

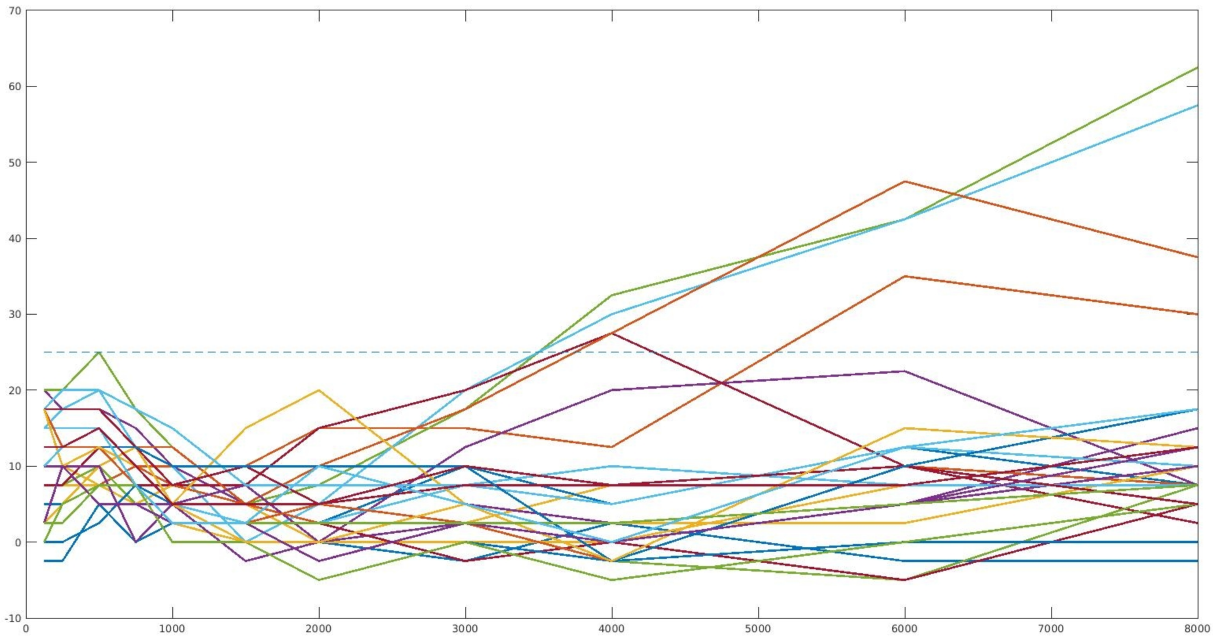


Figure 1. Averaged hearing thresholds over both ears per frequency. Audiometry is performed for both ears at 125 Hz, 250Hz, 500Hz, 1000Hz ,2000Hz ,3000Hz ,4000Hz and 8000Hz.
